# How many bits of information can be transferred between residues in a protein and how fast?

**DOI:** 10.1101/2022.01.10.475716

**Authors:** Aysima Hacisuleyman, Burak Erman

## Abstract

Time resolved Raman and infrared spectroscopy experiments show the basic features of information transfer between residues in proteins. Here, we present the theoretical basis of information transfer using a simple elastic net model and recently developed entropy transfer concept in proteins. Mutual information between two residues is a measure of communication in proteins which shows the maximum amount of information that may be transferred between two residues. However, it does not explain the actual amount of transfer nor the transfer rate of information between residues. For this, dynamic equations of the system are needed. We used the Schreiber theory of information transfer and the Gaussian network Model of proteins, together with the solution of the Langevin equation, to quantify allosteric information transfer. Results of the model are in perfect agreement with ultraviolet resonance Raman measurements. Analysis of the allosteric protein Human NAD-dependent isocitrate dehydrogenase shows that a multitude of paths contribute collectively to information transfer. While the peak values of information transferred are small relative to information content of residues, considering the estimated transfer rates, which are in the order of megabits per second, sustained transfer during the activity time-span of proteins may be significant.

## Introduction

Although mean square fluctuations of atoms in a protein are generally considered as noise due to thermal energy, they play important role in the way the protein performs its function. Each atom has a well-defined mean position dictated by the three dimensional conformation of the protein. Each atom exhibits fluctuations from its mean position. The probability distribution of these nanometer-range fluctuations is unique and time invariant. The set of all atom fluctuation distributions constitutes the unique signature of that protein. Each atom contains a specific amount of information which is transferred to other atoms as a result of correlations among fluctuations. Considering residues rather than single atoms for simplicity, the information content, *I* (Δ*R*_*i*_), of the *i*^*th*^ residue may be expressed according to the Shannon equation, *I* (Δ*R*_*i*_) = −⟨ln *p* (Δ*R*_*i*_ (*t*))⟩, where p denotes the probability, Δ*R*_*i*_ (*t*) is the instantaneous fluctuation of residue *i* from its mean position and the angular brackets denote the average. Thus it is only the time invariant distribution of fluctuations that defines the information content. When the logarithm is to base 2, the information amount is in bits. Actually, it is not the magnitude of the information contained but the way it is transferred to other residues is what determines the function. Transfer of information from one residue to another is the source of allostery in proteins. More specifically, allosteric communication or information transfer in a protein is the process in which a signal from one region of the protein, the allosteric site, is carried to another region, the active site. The signal carries a message which modulates the motion of the active site where the function of the protein is performed. Experimentally, vibrational energy flow may be observed directly by ultraviolet resonance Raman spectroscopy. In Figure 1, we reproduced the results of measurements in Cytochrome c by Mizutani group^1^, where the heme group of the protein was excited and the energy relaxation of Tryptophan 59 was monitored as a function of delay time. The heme group was excited at -5 ps. The figure shows the resulting relative energy change of Trp59 transferred from the heme group. The transferred energy is zero at zero time, increases up to a peak value and then decays and is dissipated to its environment in about 40 ps. Later experiments on Cytochrome c where three residues are mutated as Phe43Trp, Val68Trp and Leu89Trp all showed similar information transfer characteristics^2^. A large number of experiments, ultraviolet resonance Raman, NMR, and time resolved infrared spectroscopy showed the characteristic behavior of information flow in proteins^3-7^ which is also verified by molecular dynamics simulations^8, 9^. Reid et.al., ^10^ derived a scaling relation between vibrational energy transfer rates and distance between residues. In addition to direct observation of information transfer by time resolved Raman spectroscopy, evidence showing the role and importance of information transfer in biological systems is growing systematically. Balaceanu et al^11^ analyzed transfer in protein-DNA complexes and pointed to the transient nature of information transfer, an important observation which we elaborate in detail in this paper. In a recent review paper on molecular machines, Loutchko and Flechsig^12^ introduced the concept of information thermodynamics to quantify allosteric interactions. Verkhivker et al^13^ discussed the role of residues acting as drivers of the fluctuations of other residues in terms of causality in the correlations, a point which we discuss in detail in the present paper. The consequences of causality resulting in directional information flow in DNA and protein systems is pointed out by Campitelli et al^14^. Despite a relatively thorough understanding of information transfer from experiments, and molecular dynamics simulations, a molecular model that explains the amount and speed of information transfer in proteins is missing. Answers to several questions such as how much of the mutual information between two residues can be transferred from one to the other, what is the direction of transfer, which protein network parameters determine and control information transfer are not yet known. The specific purpose of the present paper is to present a simple model that explains the theoretical foundations of information transfer in proteins.

**Figure 1.**
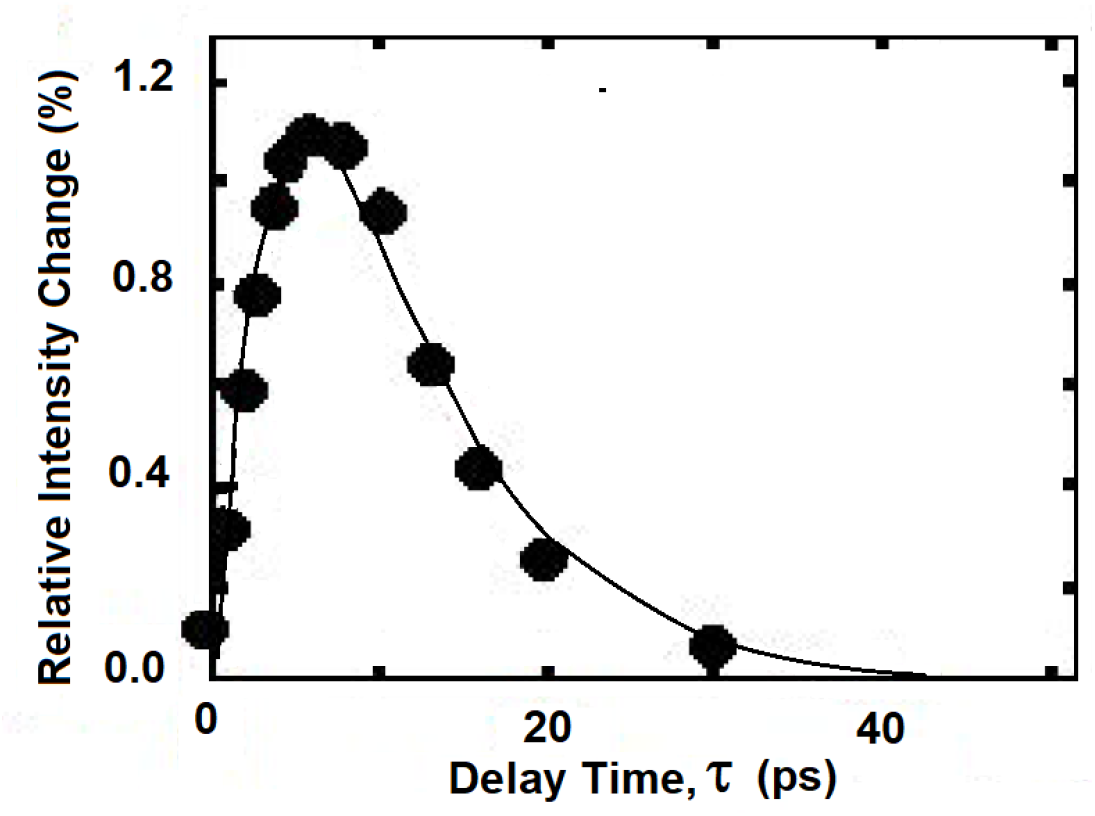
Relative vibrational intensity change of Trp59 resulting from energy transfer from Heme in Cytochrome c. Trp59 is approximately 13Å away from the heme group. The filled circles are experimental points, the curve is drawn by hand to guide the eye. The data is taken from Figure 2 of Fujii, N.; Mizuno, M.; Mizutani, Y., Direct observation of vibrational energy flow in cytochrome c. *The Journal of Physical Chemistry B* **2011**, *115* (44), 13057-13064.

In many ways, there is a strong similarity between information transfer systems used in communication technologies and allosteric proteins. Information transfer in a telephone line and an allosteric protein both have the Shannon equation in common. Basics of information mapping has been simply and clearly explained in the work of Lombardi, Holik and Vanni^15^ to which we will refer in describing information processes in proteins. Another clearly written and enlightening source on information is the book by Lesurf^16^. The figure of a general communication system given in the classic paper by Shannon^17^ and reproduced in Lombardi et al^15^ serves as a good reference also for information transfer in a protein. There will be an information source, which may be a microphone or the keyboard of a computer, an amino acid or a ligand. In telecommunication, the fluctuations in the electric current will be the source of modulation. In a protein where the source is a residue, the fluctuations of its atoms may be imagined to be sending the signal. The information source may not be confined to conformational fluctuations but as well be fluctuations in an electromagnetic field or electric charge or the atoms of a ligand docked to the protein. For simplicity we use a coarse grained model and consider the fluctuations of the alpha carbon of each amino acid. Suppose the fluctuations take only 2 equally probable values, each either +1Å or -1Å. Then the information content of this residue from the Shannon equation will be 1 bit. The effects of this pattern of fluctuations will be carried to another residue, the active site residue, or the receiver, through several paths which we will collectively call the channel. The specific pattern of the signal will constitute the message. Upon arrival at the receiver, the message will (or may) induce changes in the fluctuations of the active site residue. The information content of the receiver will be *I* (Δ*R*_*j*_) = − ⟨ln *p* (Δ*R*_*j*_ (*t*))⟩, this time *p* (Δ*R*_*j*_ (*t*)) denoting the probability of fluctuations of the active site residue *j*. This is how the information from *i* will affect *j* and lead to the function of the protein. For example, the message may slow down the fluctuations at the destination leading to a higher affinity for binding of a drug molecule to the active site residue, i.e., the receiver. However, we will not specify the ‘function’, explicitly. How a message is converted into a functional motion is similar to semantics in language which is beyond our interest in this paper.

For an illustrative example, we present in Figure 2 the molecular dynamics trajectories for the fluctuations Δ*R* of two residues of Isocitrate dehydrogenase γ subunit, chain B in 5GRH.pdb. The ordinates show the instantaneous square fluctuations of the alpha carbons of two residues LYS182 and ASP215. The right panel shows the product of their fluctuations. The product of fluctuations in the right-hand panel would show similar amounts of positive and negative values if the two residues did not communicate with each other. However, we see a strong tendency for positive values in the figure. Although the fluctuations in the left and center panels have the appearance of random noise, they contain information, as will be quantified below, and a portion of this information is shared between the two residues.

**Figure 2.**
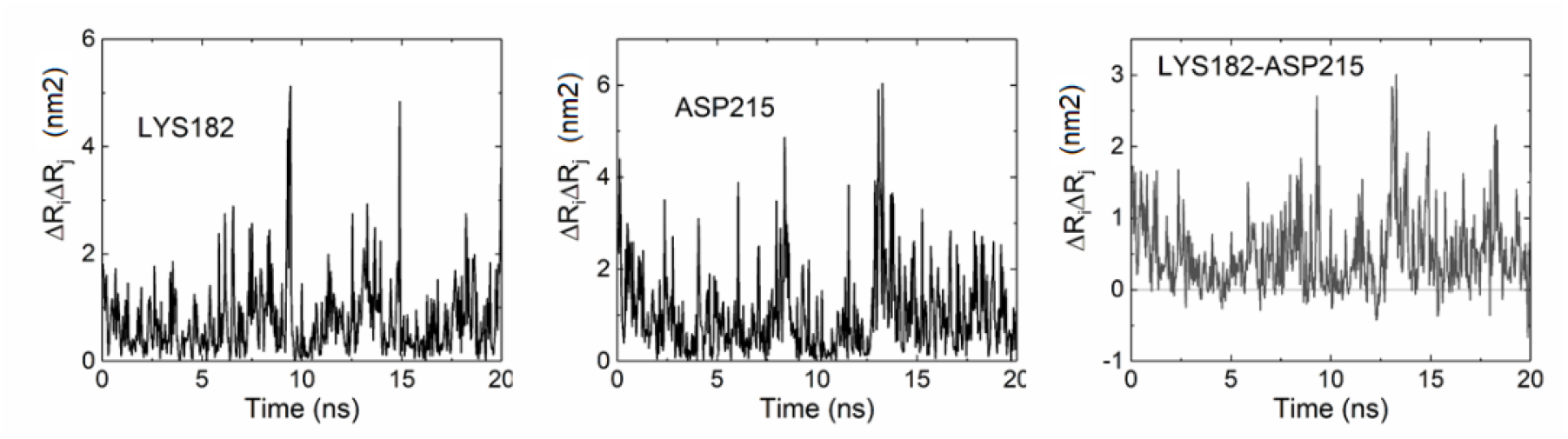
Squared fluctuations of two representative residues, LYS182 and ASP215 shown in the left-hand middle panels, respectively. Product of fluctuations is presented on the right panel. The values shown are dot products of the respective fluctuation vectors of the alpha carbons.

The relationship between the information content of residues *i* at the allosteric site, for example, and *j* at the active site and the amount shared between them may be represented by a Venn diagram^15^, Figure 3, where *I* (Δ*R*_*i*_ : Δ*R*_*j*_) is the mutual information generated at *i* and received at *j*. E is the equivocation, information generated at *i* but not received at *j*. N is the amount of information at *j* not coming from *i*, i.e., the noise that is dissipated to the rest of the protein.

**Figure 3.**
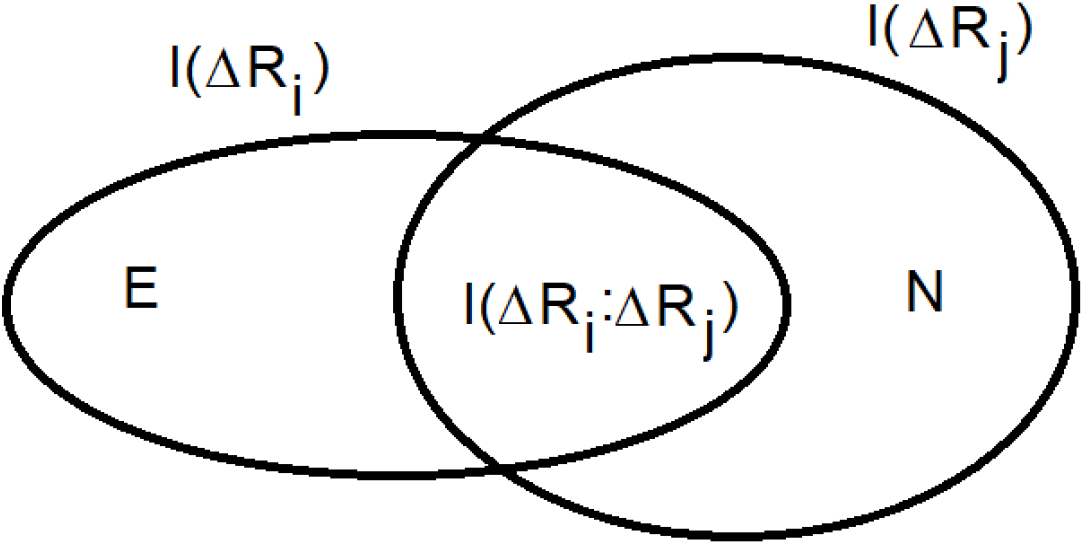
Venn diagram showing the information content and mutual information for two residues *i* and *j*. See text for description of the variables shown.

Although the Shannon equation, the Venn diagram and the concept of mutual information explain the interactions at the active and the allosteric sites, they cannot explain the transfer of information from one site to the other and how fast this takes place. In order to explain information transfer, the dynamic equations of the system are needed^18^. The theory showing the difference between mutual information and information transfer was given by Schreiber^19^, followed by the work of Kamberaj and van der Vaart^20^ and its more recent application^21, 22^ to proteins. For proteins, molecular dynamics simulations describe the dynamics of the system but the Langevin equation and harmonic interactions serve as simplifying tools^23, 24^ which were adopted in Reference ^22^ and developed further in the present paper. Here, we go beyond mutual information and consider how and how fast information is transferred from one residue to the other and vice versa. It is worth noting that mutual information is symmetric in *i* and *j*, consequently it is not possible to determine the direction of net information flow. The Schreiber theory, on the other hand, is directional and tells which way information will be transferred.

As an application of the model presented here, we investigate information transfer in the protein human NAD-dependent isocitrate dehydrogenase, 5GRH.pdb, and try to quantify the directional information flow between various residue pairs. Recently, Ding’s group presented^25^ a remarkable set of experimental data on the allosteric mechanisms of various isocitrate dehydrogenase systems which we study here in more detail from the point of view of information transfer.

## MATERIALS AND METHODS

The present analysis is based on the coarse grained model of proteins with harmonic interactions. The equations of information transfer has been presented in previous work^22^. Here we summarize and improve the theory presented in Reference ^22^. The detailed derivation of the equations are presented in the Supplementary Material section.

A residue may be visualized as a bead connected to its mean position in the crystal structure with a linear spring. In this case the probability distribution of its fluctuations will be described by a Gaussian. The information content of a residue in the Gaussian approximation is derived in the Supplementary Material as:

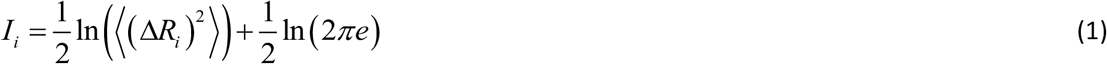

⟨(Δ*R*_*i*_)^2^⟩ is the mean-squared fluctuation obtained by taking the spring constant of the harmonic interactions as unity^26^.

Mutual information between residues *i* and *j* for harmonic interactions is derived in the Supplementary Material section as

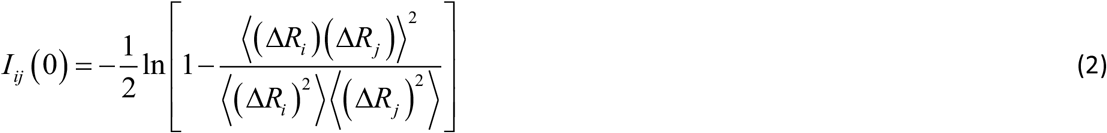

Although Equations 1 and 2 tell us how much information is contained in a given residue and how much information is shared between two residues, they do not show how much information is transferred from *i* to *j*. Transfer of information requires the knowledge of the dynamics of the system. We adopted the solution of the Langevin equation for this purpose with details discussed in the Supplementary Material section.

The transfer of information *T*_*ij*_ (*τ*) from *i* to *j* with a time delay *τ* may be written as

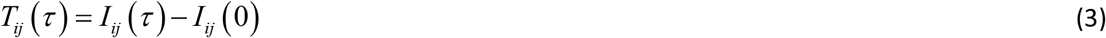

where, *I*_*ij*_ (0) is the time independent mutual information given by Eq 2 and *I*_*ij*_ (*τ*) is the time dependent mutual information given by Eq. 14 of the Supplementary Material section as:

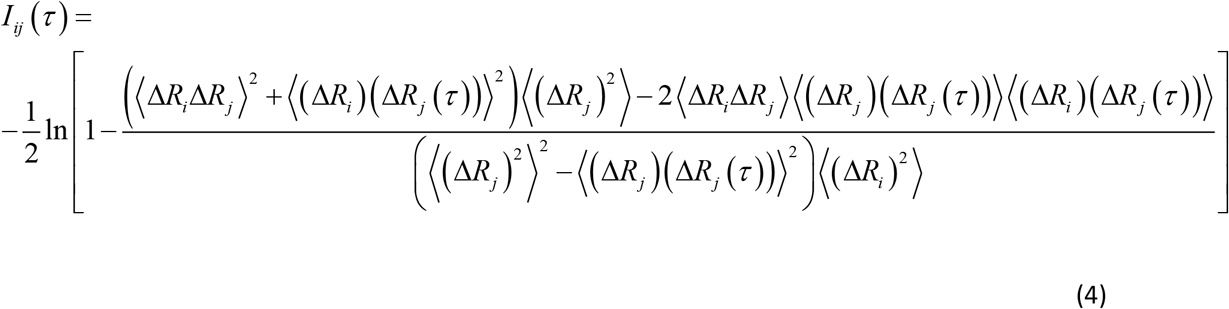

where,

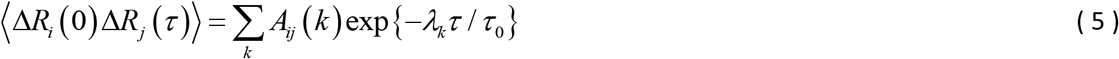

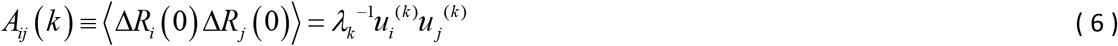

with *λ*_*k*_ being the *k*^*th*^ eigenvalue and *u*_*i*_^(*k*)^ being the *i*^*th*^ component of the *k*^*th*^ eigenvector of the connectivity matrix of the GNM, Γ. The characteristic time *τ*_0_ is defined as a universal time constant first given by Ben Avraham as a universal vibrational spectral property of proteins^27^ and later shown^24^ to be around 6 ps.

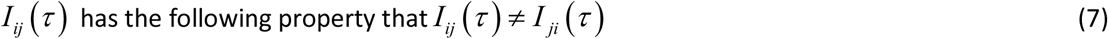

In all calculations, we divide the information transfer expression, Eq. 3, by ln 2 which expresses information transfer in bits.

Information originating at i and arriving at j at time *τ* is not equal to information originating at j and arriving at i at *τ*. The difference

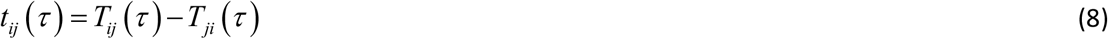

between them is the net information transfer from i to j, observed at time *τ*. Equation 8 specifies the directionality of information flow.

## Results and discussion

Human NAD-dependent isocitrate dehydrogenase, Protein Data Bank code 5GRH.pdb, catalyzes the decarboxylation of isocitrate into α-ketoglutarate in the Krebs cycle. It is allosterically activated by the binding of citrate and ADP. The protein is a dimer of α and γ subunits. The citrate, ADP, and Magnesium bind adjacent to each other at the allosteric site on the γ subunit. Binding of the citrate at the allosteric site of the γ subunit induces conformational changes at the allosteric site, which are transmitted to the active site on the α subunit through the heterodimer interface. The information coming from the γ subunit leads to stabilization of the ligand isocitrate, (ICT) binding at the active site on the α subunit resulting in the activation of the enzyme. Citrate interacts with LYS76, ASN78, THR81, SER91, ASN93, ARG97, ARG128, TYR135, LYS175, ARG272 of the allosteric site of the γ subunit. Magnesium interacts with ASN78 and ARG272. The active site of the α subunit consists of two amino acids, ASP230 and ASP234. These two residues are in direct contact with the VAL214-SER223 helix of the γ subunit at the interface.

The heterodimer is shown in Figure 4. The α subunit is on the left. The active site residues ASP230 and ASP234 in the α subunit are shown as black spheres. ASP215 of the γ subunit shown as a gray sphere is at the interface. It is located on a helix which neighbors the active sites residues of the α subunit. The allosteric site residues LYS175 and ARG128 are shown on the right side of the γ subunit as gray spheres. As will be shown below, binding of the ligand at LYS175 sends information to ARG128 which through a cascade of pathways sends information to ASP215 which acts on the two active site residues of the α subunit through the interface, resulting in the activation of the enzyme. This communication, which originates at LYS175 results in information transfer through the cascade of pathways which all converge around the residues VAL232 and finally at ASP215, as will be explained below in detail.

**Figure 4.**
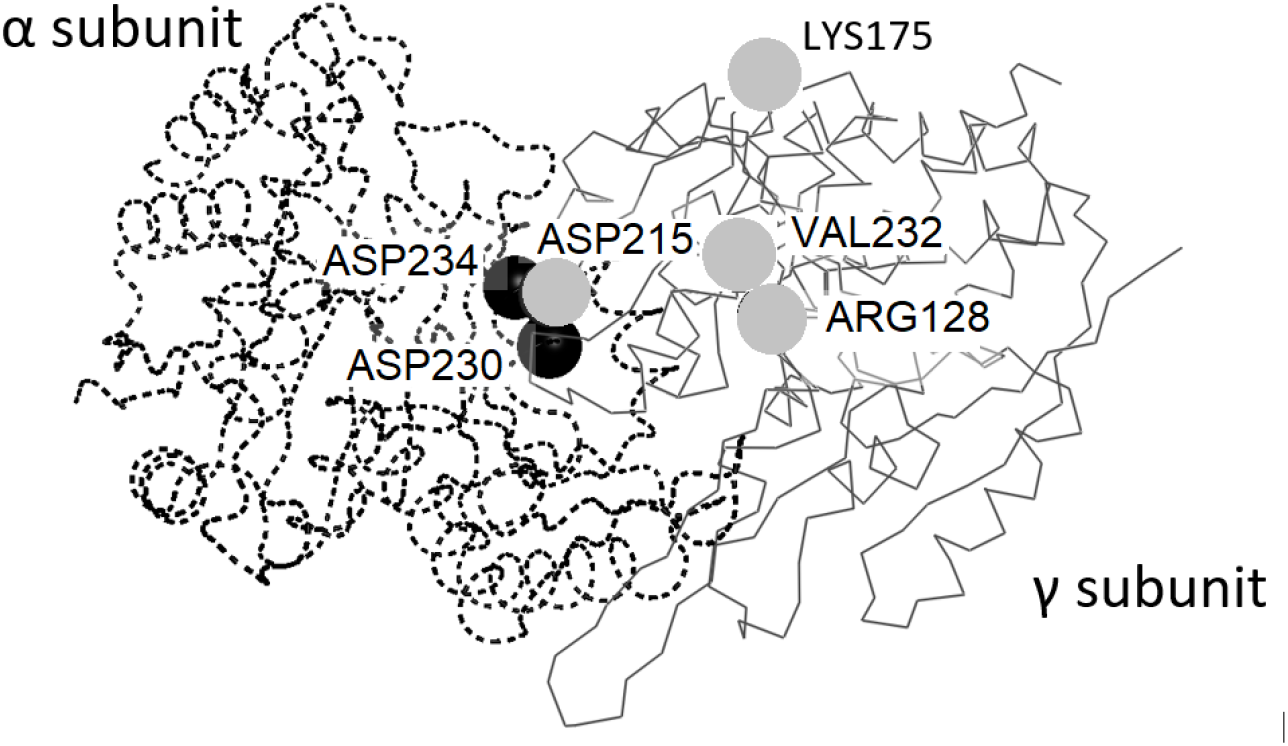
The α and γ subunits of 5GRH.pdb. Information flows from LYS175 to residues around ARG128 and its spatial neighbor VAL232. Information is then distributed into several paths converging at a helix that contains ASP215. The active site residues of the α subunit, ASP230 and ASP234, are shown as black spheres.

The values of net information transfer *t*_*ij*_ vary in the range -0.04 to 0.04. The overall map of *t*_*ij*_ in the system is presented in Figure 5 for the γ subunit. The left panel shows all positive values of *t*_*ij*_. In the right panel, information transfer values of *t*_*ij*_ greater than 0.4% are shown. The abscissae in both panels show the residues from which information originates. The ordinates show the residues to which information is transferred. The central vertical strip spanning the abscissa values from 128 to 223 on the left panel defines an information transfer block in the protein showing the residues from which information flows. These residues form a reservoir of information. Several horizontal streaks are observed in this block, their ordinate values indicating the residues to which information flows. The region circled on the right panel indicates significant information flow into residues 216-233 (ordinate values). The terminal segment of this range, residues 215-221, belongs to a helix that neighbors the two active site residues, ASP230 and ASP234, of the α subunit. We can state that all information coming from the reservoir is transferred to the active sites of the α subunit through this helix.

**Figure 5.**
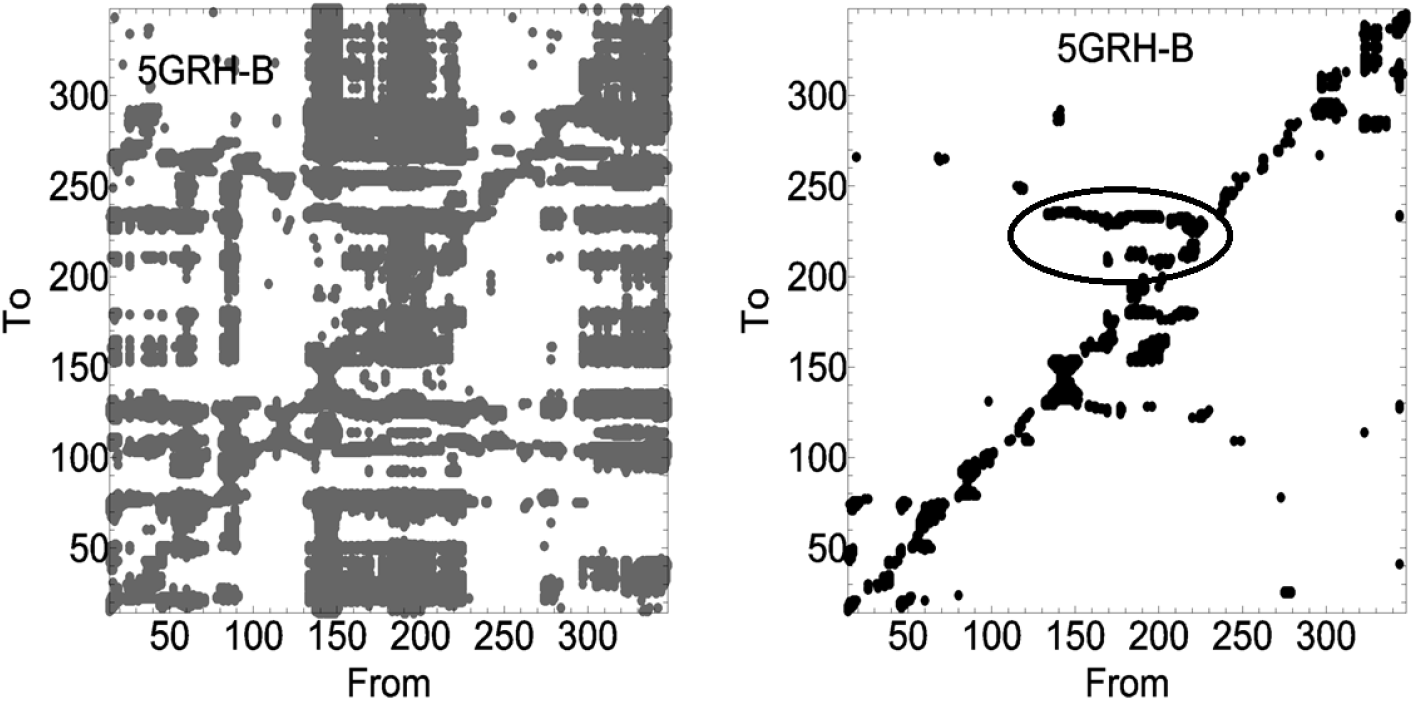
Net information transfer *t*_*ij*_ in the protein. Left panel shows all positive values of *t*_*ij*_. Maximum *t*_*ij*_ value is 0.04. Right panel shows only the values above 0.004. The encircled region on the right panel indicates the residues with significant allosteric activity.

The residues that are important in carrying the information also appear in the mutual information diagram, Figure 6, where the horizontal streak encircled in the right panel of Figure 5 is also visible here. The mutual information values vary between 0 and 0.7 and are in the order of ten to twenty times higher than information transfer values. According to the data, maximum mutual information is between ARG128, LYS175 and the interface helix 215-221. These are the residues which appear most frequently in Figure 5. Their frequencies of occurrence are presented in Figure 7.

**Figure 6.**
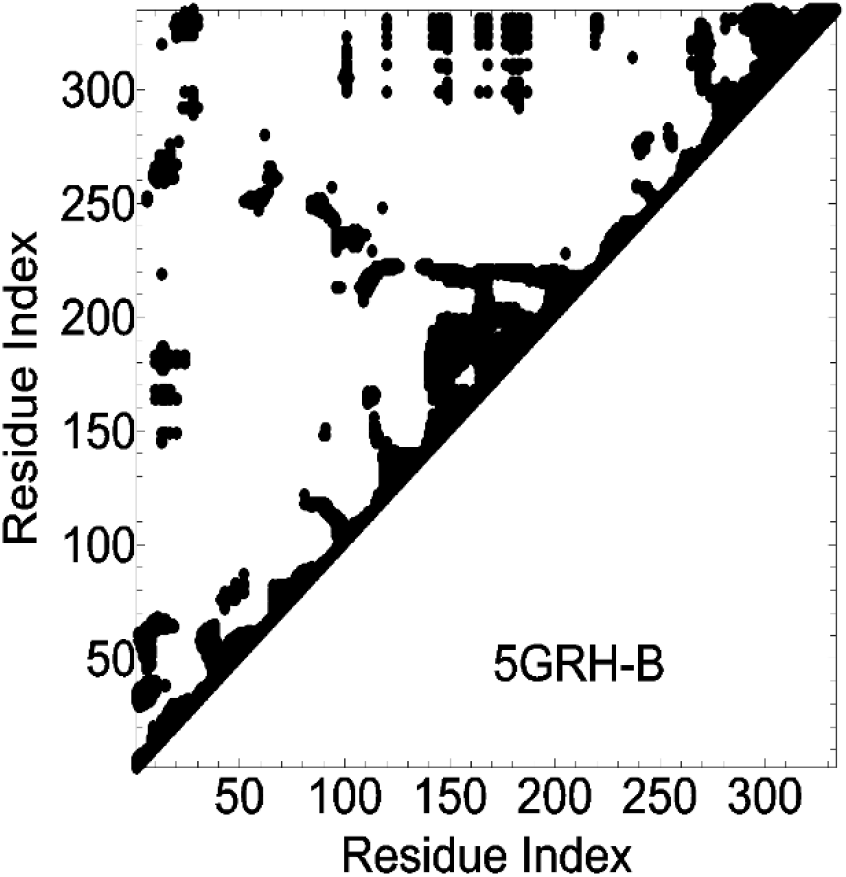
Mutual information as a function of residue index. Only half of the data is represented since the function is symmetric.

**Figure 7.**
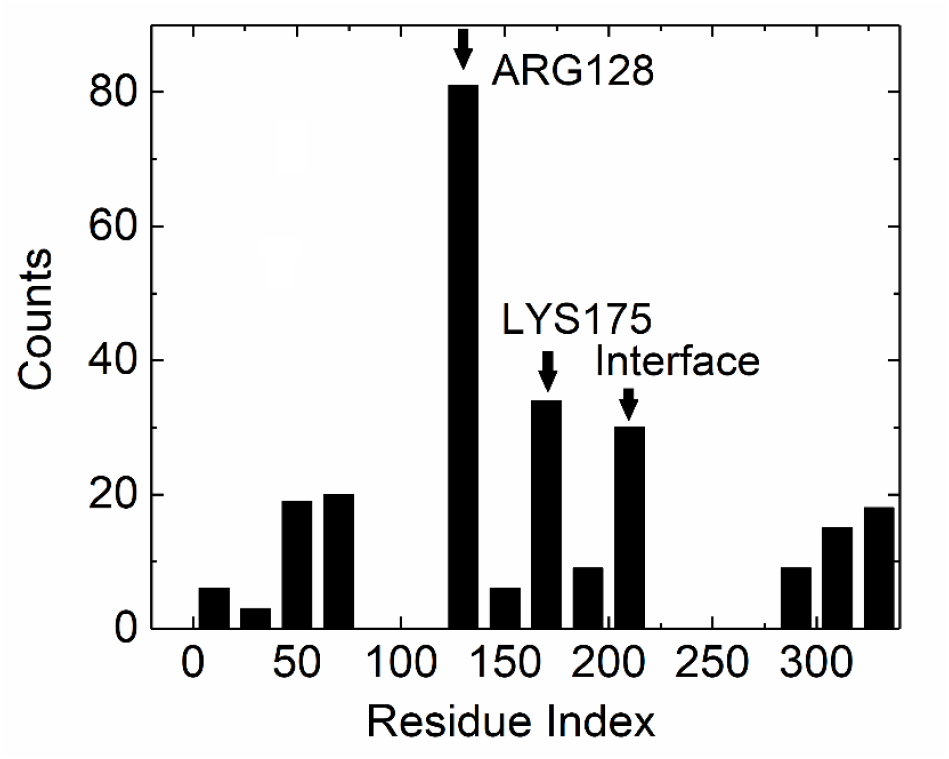
Number of occurrence (ordinate values) of residues with top 10% of values of mutual information.

### Information transfer between specific residue pairs

Information transfer between residues that are involved in allosteric activity of isocitrate dehydrogenase are presented in Figure 8. The ordinate values show the amount of information transferred, abscissa values denote the time lag of observation, i.e., the observation time *τ* at j when the information originates at i at time zero. At *τ* = 0 information transferred to j is zero because it takes a finite time for information to travel, the characteristic time being around 6 ps as suggested earlier^27^. This is built into the Langevin equation whose solution is given by Eq. 5. The peak information is observed around this characteristic time. At longer times, the information starts to decay due to noise in the system. It should be noted that the fluctuations at j are also interacting with several other residues, in fact with all other residues of the protein to different extents, and this constitutes the noise. At infinite time, no information can be observed at j. The curves in Figure 8 clearly show the causality of information transfer. The solid curves show transfer from i to j, the dot-dashed curve shows from j to i. The larger ordinate values show the dominant direction of information flow. The first residue number in the label in each figure represents i, the second is j.

**Figure 8.**
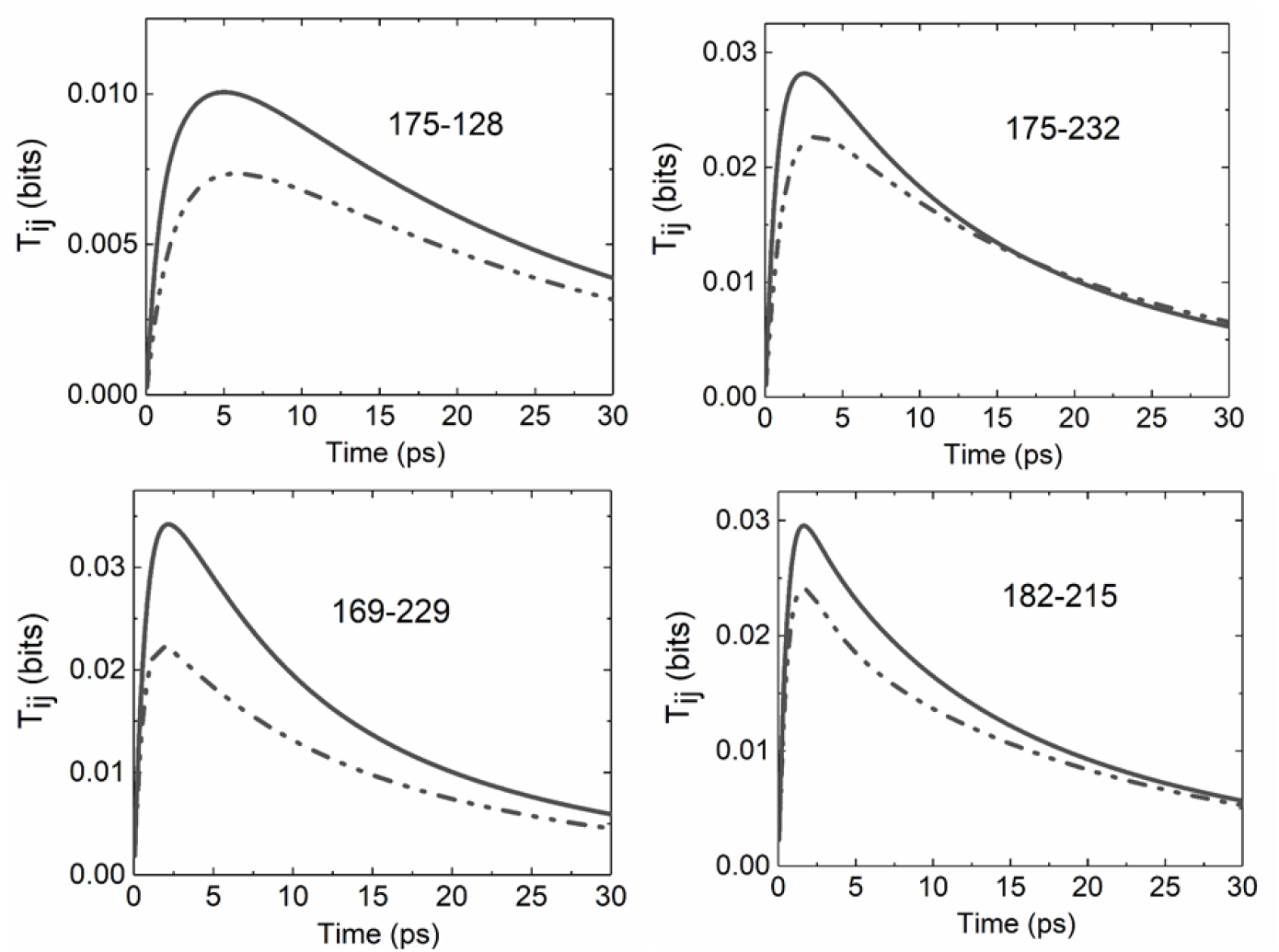
Information transfer curves for some pairs. Solid lines are in the order marked in each figure. Dot dashed lines are in the opposite direction

#### LYS175-ARG128 pair

Information transfer between these two residues is presented in the upper left panel of Figure 8. Their alpha carbons are at a distance of 18.3Å. They communicate with each other, information going from LYS175 on which the citrate binds to ARG128. The information content of LYS175 and ARG128, calculated with the Shannon equation, are 4.75 and 4.29 Bits respectively. However, the mutual information between the two, calculated with Eq. 2, is much smaller, 0.013 Bits, showing that most of the information from LYS175 is lost as equivocation, i.e., does not affect ARG128, and most of the information content of ARG128 is lost as noise concerning their interaction. Peak information transfer from LYS175 is observed at a time delay of 5 ps. And the observed amplitude is 0.010 Bits, which is approximately 77% of the mutual information. If, as a coarse approximation, we assume that all of the information is concentrated at the peak, then dividing the peak information by the peak time gives a measure of the speed of propagation, which is 2.01 Mbits/sec. This is the maximum rate at which information may be transferred from LYS175 to ARG128. In the opposite direction, transfer is slower with a peak time of 5.75 ps. The peak transfer is 0.007 Bits with a peak rate of 1.28 Mbits/sec.

Values for the other pairs that are closely related to allosteric communication are presented in Table 1.

**Table 1.** Information transfer data for residues involved in allosteric communication

| Residue Pair | C<sup>α</sup>-C<sup>α</sup> Distance Å | Information content (Bits) | Mutual Information (Bits) | Peak Time (ps) | Peak Transfer (Bits) | Peak rate (Mbits/sec) |
| --- | --- | --- | --- | --- | --- | --- |
| LYS175-ARG128 | 18.3 | 4.75 | 0.013 | 5.00 | 0.011 | 2.01 |
|  |  | 4.29 |  | 5.75 | 0.007 | 1.28 |
| LYS175-VAL232 | 13.7 | 4.75 | 0.054 | 2.60 | 0.028 | 10.83 |
|  |  | 4.33 |  | 3.40 | 0.023 | 6.66 |
| ALA169-ASP229 | 8,8 | 4.63 | 0.080 | 2.20 | 0.034 | 15.55 |
|  |  | 4.48 |  | 1.85 | 0.022 | 12.02 |
| LYS182-ASP215 | 8.0 | 4.72 | 0.131 | 1.70 | 0.030 | 17.39 |
|  |  | 4.86 |  | 1.70 | 0.024 | 14.15 |

#### LYS175-VAL232

Binding of citrate perturbs LYS175, and information from this and several other residues are transferred to and converge at VAL232. The information transfer is presented in the upper right panel of Figure 8. VAL232 is spatially close to interface residues and affects the motions of the two active site residues of the α subunit. The peak value of transfer is larger than the previous case.

#### ALA169-ASP229

ALA169, which is spatially close to LYS175 is another residue through which information is sent to the information reservoir. ASP229 is a representative residue of the reservoir. Information flow from ALA169 to ASP229, whose alpha carbons are 8.8Å apart is shown in the lower left panel of Figure 8. The peak value is higher than the previous two discussed here, and the causality effect is high as may be seen from the difference between the solid and the dot dashed curves.

#### LYS182-ASP215

LYS182 is another residue of the reservoir from which information is sent to the active residues. Here, LYS182 sends information directly to ASP215 is at the interface and contacts the active site residues of the α subunit. The peak value and the degree of causality is high.

The mutual information and information transfer values shown in Table 1 are considerably smaller than the information content of the residues. Then, one is confronted with the question of how allosteric communication can be effective with such small transfer values? There may be two answers to this question: Firstly, there are several pairs that transfer information to the interface as may be seen from the left panel of Figure 5. Since information transfer is additive, the sum of the transfers is expected to be highly effective. Secondly, information transfer rates are high and continuous flow of information between residues may be significant.

## Conclusion

We used the solution of the Langevin equation for the Gaussian Network Model and Schreiber’s theory of entropy transfer to describe information transfer in proteins. The quantitative time decay curves of information transfer that are obtained agree perfectly with results of ultraviolet resonance Raman spectroscopy experiments. The curves peak around 5 ps and decay to zero in 30-40 ps. The time required for information to reach a residue is around 5 ps as given by both experiment and theory. As observed in experiments^2^, amplitude of information transfer decreases with distance between residues. Peak time increases with pair distance, also in agreement with experiments.

The dominant factor in information transfer is the individual and joint probability distributions of the fluctuations. In Raman experiments^1^ on Cytochrome-c the heme is pumped with a laser to a higher energy level which then propagates to the neighboring Tryptophan. The jump in heme energy is a fluctuation, resulting from the perturbation by the laser source and is in the form of a Dirac spike. The theory, on the other hand, is derived in the absence of perturbation, simply using the joint probabilities of equilibrium fluctuations for transfer. Although a strong perturbation is expected to induce stronger correlations and hence stronger transfer, a recent study^23^ showed that effects induced by perturbations ride on the existing equilibrium correlations. This observation was obtained by the solution of the Langevin equation in the absence of perturbations.

Finally, the present simple Gaussian model can help explain the mechanism of the amount and speed of information transfer between two residues as well as the direction of the transfer. The agreement of theory and experimental time scales is encouraging for future more detailed work in this area.

## Data Availability

The data that support the findings of this study are available from the corresponding author upon reasonable request.

