## Supplemental File for "How many bits of information can be transferred between residues in a protein and how fast?"


In this section, we give detailed derivations of the information transfer equations presented in the main manuscript.

Information transfer is a dynamic process and the solution of the equation of motion is needed. Here, we adopt the coarse-grained Langevin dynamics model of proteins[1, 2]

$$\varsigma \Delta R' + \Gamma \Delta R = F \quad (1)$$

where, $\varsigma$ is the effective friction coefficient, $\Delta R$ is the vector that represents the instantaneous fluctuation of residues, the prime is the time derivative, $F$ is the external force acting on the residues arising from random noise. In the coarse graining approximation we assume that the atoms of each residue is collapsed on its alpha carbon. The vectors shown in Eq. 1 have n elements which is the number of alpha carbons in the protein. $\Gamma$ in Eq. 1 is the spring constant matrix, defined in Reference 1. The choice of $\Gamma$ in this form is a result of the assumption of a protein with harmonic interactions and leads to the Gaussian Network Model.

Solution of the Langevin equation leads to time delayed correlations derived in references [1, 2]:

$$\left\langle \Delta R_i(0) \Delta R_j(\tau) \right\rangle = \sum_k A_{ij}(k) \exp\left\{ -\lambda_k \tau / \tau_0 \right\} \quad (2)$$

Where,

$$A_{ij}(k) = \lambda_k^{-1} u_i^{(k)} u_j^{(k)} \quad (3)$$

with $\lambda_k$ being the k$^{th}$ eigenvalue and $u_i^{(k)}$ being the i$^{th}$ component of the k$^{th}$ eigenvector of $\Gamma$.

The relationship between $\Gamma$ and equilibrium fluctuations is given as[3]

$$\left\langle \Delta R_i(0) \cdot \Delta R_j(0) \right\rangle = \left( \Gamma^{-1} \right)_{ij} \quad (4)$$

In the derivation of Eq. 4 a factor of $k_B T / \gamma$, where $k_B$ is the Boltzmann constant, $T$ the temperature, and $\gamma$ the spring constant of elastic interactions is set to unity since we are interested in information transfer between residues.

The probability distribution of a Gaussian of n variables, $\Delta R=\left[\Delta R_1,\Delta R_2,\Delta R_3,\cdots,n\right]$ is

$$p\left(\Delta R\right)=\frac{1}{\sqrt{\left(2\pi\right)^n\det\Gamma^{-1}}}\exp\left(-\frac{1}{2}\left(\Delta R\right)^T\Gamma\left(\Delta R\right)\right) \tag{5}$$

Using Eq. 5, one obtains different orders of joint probabilities and information expressions. For example, the information content $I_i=-\left\langle \ln p\left(\Delta R_i\left(t\right)\right)\right\rangle$ upon use of Eq. 5 is obtained as

$$I_i=\frac{1}{2}\ln\left(\left\langle\left(\Delta R_i\right)^2\right\rangle\right)+\frac{1}{2}\ln\left(2\pi e\right) \tag{6}$$

For pairwise dependence, Eq. 5 gives

$$\left\langle \ln p\ \left(\Delta R_i,\Delta R_j\right)\right\rangle=-\frac{1}{2}\ln\left(\left\langle\left(\Delta R_i\right)^2\right\rangle\left\langle\left(\Delta R_j\right)^2\right\rangle-\left\langle\left(\Delta R_i\right)\left(\Delta R_j\right)\right\rangle^2\right)-\ln\left(2\pi e\right) \tag{7}$$

And mutual information defined in general as

$$I\left(\Delta R_i,\Delta R_j\right)=\left\langle \ln\frac{p\left(\Delta R_i,\Delta R_j\right)}{p\left(\Delta R_i\right)p\left(\Delta R_j\right)}\right\rangle=\left\langle \ln p\left(\Delta R_i,\Delta R_j\right)\right\rangle-\left\langle \ln p\left(\Delta R_i\right)\right\rangle-\left\langle \ln p\left(\Delta R_j\right)\right\rangle \tag{8}$$

Gives for the Gaussian approximation:

$$I\left(\Delta R_i,\Delta R_j\right)=-\frac{1}{2}\ln\left[1-\frac{\left\langle\left(\Delta R_i\right)\left(\Delta R_j\right)\right\rangle^2}{\left\langle\left(\Delta R_i\right)^2\right\rangle\left\langle\left(\Delta R_j\right)^2\right\rangle}\right] \tag{9}$$

Equations (6) and (9) gives the information content in a residue and mutual information of two residues, but cannot give how information is transferred from one residue to another. For this, we need the information transfer model given by Schreiber[4] which reads in its original form as

$$\begin{aligned}T_{i\to j}\left(\tau\right)=&-\left\langle \ln p\ \left(\Delta R_j\left(0\right),\Delta R_j\left(\tau\right)\right)\right\rangle+\left\langle \ln p\ \left(\Delta R_i\left(0\right),\Delta R_j\left(0\right),\Delta R_j\left(\tau\right)\right)\right\rangle\\&+\left\langle \ln p\ \left(\Delta R_j\left(0\right)\right)\right\rangle-\left\langle \ln p\ \left(\Delta R_i\left(0\right),\Delta R_j\left(0\right)\right)\right\rangle\end{aligned} \tag{10}$$

In the Gaussian approximation, Eq. 10 is shown by Hacisuleyman and Erman[5] to reduce to

$$
\begin{aligned}
T_{i\to j}(\tau) &= \frac{1}{2}\ln\left(\left\langle\left(\Delta R_j(0)\right)^2\right\rangle^2 - \left\langle\left(\Delta R_j(0)\right)\left(\Delta R_j(\tau)\right)\right\rangle^2\right) \\
&-\frac{1}{2}\ln\Big[\ \left\langle\left(\Delta R_i(0)\right)^2\right\rangle\left\langle\left(\Delta R_j(0)\right)^2\right\rangle^2 \\
&+2\left\langle\Delta R_i(0)\Delta R_j(0)\right\rangle\left\langle\Delta R_j(0)\Delta R_j(\tau)\right\rangle\left\langle\Delta R_i(0)\Delta R_j(\tau)\right\rangle \\
&-\left\{\left\langle\Delta R_i(0)\Delta R_j(\tau)\right\rangle^2 + \left\langle\Delta R_i(0)\Delta R_j(0)\right\rangle^2\right\}\left\langle\left(\Delta R_j(0)\right)^2\right\rangle \\
&-\left\langle\Delta R_j(0)\Delta R_j(\tau)\right\rangle^2\left\langle\left(\Delta R_i(0)\right)^2\right\rangle\ \Big] \\
&-\frac{1}{2}\ln\left[\left\langle\left(\Delta R_j(0)\right)^2\right\rangle\right] \\
&+\frac{1}{2}\ln\left(\left\langle\left(\Delta R_i(0)\right)^2\right\rangle\left\langle\left(\Delta R_j(0)\right)^2\right\rangle - \left\langle\left(\Delta R_i(0)\right)\left(\Delta R_j(0)\right)\right\rangle^2\right)
\end{aligned}
\tag{11}
$$

Equation (11) can be rearranged into a simpler form by the following manipulations:

$$
\begin{aligned}
T_{i\to j}(\tau) &= -\frac{1}{2}\ln\left[1-\frac{\left(\left\langle\Delta R_i\Delta R_j\right\rangle^2 + \left\langle\left(\Delta R_i\right)\left(\Delta R_j(\tau)\right)\right\rangle^2\right)\left\langle\left(\Delta R_j\right)^2\right\rangle - 2\left\langle\Delta R_i\Delta R_j\right\rangle\left\langle\left(\Delta R_j\right)\left(\Delta R_j(\tau)\right)\right\rangle\left\langle\left(\Delta R_i\right)\left(\Delta R_j(\tau)\right)\right\rangle}{\left(\left\langle\left(\Delta R_j\right)^2\right\rangle^2 - \left\langle\left(\Delta R_j\right)\left(\Delta R_j(\tau)\right)\right\rangle^2\right)\left\langle\left(\Delta R_i\right)^2\right\rangle}\right] \\
&+\frac{1}{2}\ln\left[1-\frac{\left\langle\Delta R_i\Delta R_j\right\rangle^2}{\left\langle\left(\Delta R_i\right)^2\right\rangle\left\langle\left(\Delta R_j\right)^2\right\rangle}\right]
\end{aligned}
\tag{12}
$$

The last term is the negative of mutual information. It is symmetric in i and j and independent of time. The first term depends on time. Transfer of information equates to the mutual information term at time 0 and infinity. We may represent it as time dependent mutual information and write Eq. 12 as

$$
T_{i\to j}(\tau) = I_{ij}(\tau) - I_{ij}(0) \tag{13}
$$

Where time dependent mutual information $I_{ij}(\tau)$ is

$$I_{ij}\left(\tau\right)=$$

$$-\frac{1}{2}\ln\left[1-\frac{\left(\left\langle \Delta R_i \Delta R_j\right\rangle^2+\left\langle\left(\Delta R_i\right)\left(\Delta R_j\left(\tau\right)\right)\right\rangle^2\right)\left\langle\left(\Delta R_j\right)^2\right\rangle-2\left\langle \Delta R_i \Delta R_j\right\rangle\left\langle\left(\Delta R_j\right)\left(\Delta R_j\left(\tau\right)\right)\right\rangle\left\langle\left(\Delta R_i\right)\left(\Delta R_j\left(\tau\right)\right)\right\rangle}{\left(\left\langle\left(\Delta R_j\right)^2\right\rangle^2-\left\langle\left(\Delta R_j\right)\left(\Delta R_j\left(\tau\right)\right)\right\rangle^2\right)\left\langle\left(\Delta R_i\right)^2\right\rangle}\right] \quad (14)$$


[1] T. Haliloglu, I. Bahar, and B. Erman, Physical Review Letters **79** (1997) 3090.
[2] A. Hacisuleyman *et al.*, The Journal of Physical Chemistry B **125** (2021) 729.
[3] I. Bahar, A. R. Atilgan, and B. Erman, Fold Des **2** (1997) 173.
[4] T. Schreiber, Physical Review Letters **85** (2000) 461.
[5] A. Hacisuleyman, and B. Erman, Proteins: Structure, Function, and Bioinformatics (2017)
